# Behavior-relevant periodized neural representation of acoustic but not tactile rhythm in humans

**DOI:** 10.1101/2024.12.09.627475

**Authors:** C Lenoir, T Lenc, R Polak, S Nozaradan

**Affiliations:** Institute of Neuroscience (IONS), UCLouvain, Brussels, 1200, Belgium; Basque Center on Cognition, Brain and Language (BCBL), Donostia-San Sebastian, Spain; RITMO Centre for Interdisciplinary Studies in Rhythm, Time and Motion, University of Oslo, Norway; Department of Musicology, University of Oslo, Norway; MARCS Institute for Brain, Behaviour and Development, Western Sydney University, Sydney, 2751, Australia; International Laboratory for Brain, Music and Sound Research (BRAMS), Montreal, QC H3C 3J7, Canada

## Abstract

Music makes people move. This human propensity to coordinate movement with the rhythm of music requires multiscale temporal integration, allowing fast sensory events composing rhythmic input to be mapped onto slower internal templates such as periodic beats. Relatedly, beat perception has been shown to involve a selective enhancement of the beat periodicities in the neural response to rhythm. However, the extent to which this ability to move to the beat, and the related *periodized* neural representation, are shared across the senses beyond audition remains unknown. Here we addressed this question using acoustic and tactile rhythms, while recording separately the electroencephalography (EEG) responses and finger tapping to these rhythms in healthy volunteers. Consistent with previous studies, EEG responses to the acoustic rhythm featured significant enhancement of the beat, and this periodized neural representation was specifically concentrated within a low-frequency range below 15 Hz. In contrast, the same rhythm conveyed with tactile inputs elicited responses over a broader frequency range, up to 25Hz, with no significant periodization, and resulted in less stable tapping. Together, these findings indicate that low-frequency neural activity preferentially supports behavior-relevant internal representation of rhythmic input. However, this neural representation is not necessarily shared across the senses, as well as the ability to move to the beat, corroborating multimodal differences in beat perception. This low-frequency neural representation may thus reflect a process of multiscale temporal integration allowing the auditory system to go beyond mere tracking of onset timing and support higher-level internal representation and motor entrainment to rhythm.

**Significance statement:** Integrating the fast sensory events composing music into slower temporal units is a cornerstone of beat perception and social interaction through music. The current study shows that this ability critically relies on brain activity concentrated in a lower frequency range – below the recurrence of sensory events – in response to acoustic rhythm. In contrast, when the rhythm is conveyed through touch, brain responses comparatively exhibit higher frequency activity corresponding to the faithful tracking of each individual events of the tactile rhythm. Most importantly, these auditory-specific slow fluctuations feature a periodization of rhythmic inputs, compatible with behavior. This higher-level neural processing of rhythmic input could thus reflect internal representations of the beat that are not necessarily shared across sensory modalities, in line with the concept of auditory dominance in temporal event perception and motor entrainment to rhythm. The current study thus opens a promising avenue to gain fundamental knowledge on high-level multimodal perception and motor entrainment processes specific to humans.

## Introduction

The propensity of humans to coordinate movement with musical rhythm is attributed to a close coupling of the auditory and motor systems (Chen et al., 2008; Patel and Iversen, 2014; Patel 2024). This audiomotor coordination is thought to rely on the auditory system’s ability to integrate information at multiple timescales (Teng et al., 2016). Such multiscale temporal integration is arguably essential for human-specific social interactions such as music and dance (Patel et al., 2005; Zatorre et al., 2007), and speech (Norman-Haignere et al., 2022; Arnal et al., 2015). Specifically, this multiscale temporal scaffolding is crucial for organizing fast time intervals composing rhythmic input into slower, behavior-relevant templates. Once mapped onto the rhythmic input, these internal templates can be experienced as periodic beats and can be used to guide motor coordination with others and the music (London, 2012; Large and Snyder 2009; Honing and Bouwer 2018).

Musical beat usually refers to an internal representation consisting of recurring periods, or periodic pulses, mapped onto the rhythmic input (Honing and Bouwer, 2018; London et al., 2017; Lenc et al., 2021). How this internal periodic template is mapped onto complex sensory inputs such as music is far from trivial, especially since the beat periodicities are often not prominent in the physical structure of the input (London, 2012; Honing and Bouwer, 2018; London et al., 2017; Lenc et al., 2021). In other words, this template matching shows some invariance to the physical structure of the input in which undoubtedly makes musical beat a higher-level perceptual process. (Nozaradan et al., 2017; Lenc et al., 2021; 2024). Uncovering these processes thus offers key insights into high-level perception and motor entrainment.

Recent studies capturing human brain activity using electroencephalography (EEG) revealed that neural representation of the rhythmic inputs exhibit selectively emphasized beat periodicities, regardless of their prominence in the input (Nozaradan et al., 2017; Lenc et al., 2020; Lenc et al., 2023). Importantly, such *periodized* neural representation seems functionally relevant, as the enhanced beat-related periodicities in neural activity correspond to those preferentially expressed through body movements (Lenc et al. 2018; Nozaradan et al. 2012; 2018).

However, whether the neural enhancement of beat representation and the ability to move to it are auditory-specific or generalize beyond audition remains unknown. Synchronization to rhythm is generally recognized to be facilitated with audition, as compared to vision or touch (Hove et al., 2013; Repp&Penel 2004; Gilmore and Russo, 2021). Yet, recent studies challenged this view by showing synchronization performance close to the one typically found with audition, when the rhythmic input was tuned to match the sensitivity of the alternative sensory system. Specifically, visuomotor synchronization was considerably improved in response to moving rather than static visual rhythms, thus likely benefiting from the temporal sensitivity of vision to biological (Su and Pöppel, 2012;), and realistic trajectories (Hove et al., 2010; Gan et al., 2015; Gu et al., 2020).

Here, we addressed the question of cross-sensory commonalities and specificities in beat processing, by separately recording behavioral and EEG responses to an acoustic and tactile version of the same rhythm. Since sounds and vibrations share similar physical attributes and often concomitantly occur in musical contexts (Merchel & Altinsoy 2018; Reybrouck et al. 2019), it could be hypothesized to find similar neural emphasis on the beat to those input (Schurmann et al., 2006; Rahman et al., 2020). Yet, recent studies on sensorimotor synchronization with tactile vs. acoustic rhythms, and corresponding EEG activity (Brochard et al., 2008; Tranchant et al., 2017; Gilmore and Russo, 2021), showed discrepancies between these modalities. These differences could have been driven by the use of stimuli not specifically adjusted for the somatosensory system (i.e., the body part stimulated, the frequency sensitivity of mechanoreceptors). Here, we thus moved a critical step forward by investigating beat processing using stimuli fine-tuned to match the sensitivity of each sense, while controlling for lower-level confounds using rhythms whose physical structure does not feature prominent beat periodicities.

## Materials and Methods

### Participants

Forty-five healthy volunteers (32 women; mean age = 23.5 years; SD = 3.4) took part in the study. All participants reported normal hearing, no alteration of cutaneous sensitivity at the level of the fingers and no history or presence of neurological or psychiatric disorders. Most of the participants reported having grown up in countries from a Western culture (42 out of 45). They reported a range of musical training (mean = 1.6 years of formal music training, e.g., music lessons; SD = 3.0; range 0-14 years), with 8 of them self-identifying as “amateur musicians” while the other 37 participants defined themselves as “non-musicians”.

Participants were randomly assigned to one of the three groups corresponding to three different blocks orders, i.e. group 1 : first tactile – second acoustic – third tactile (n = 15; mean age = 25.9 years; SD = 4.3; 12 women), group 2: first tactile – second tactile – third acoustic (n = 15; mean age = 25.4 years; SD = 2.3; 12 women), and group 3: first acoustic – second tactile – third tactile (n = 15; mean age = 24.7 years; SD = 3.4; 8 women). Participants provided written consent after being informed about the experimental procedures. All procedures were approved by the local ethical committee *Comité Hospitalo-facultaire de l’UCLouvain* (protocol number B403201938913).

### Rhythmic sequences

Participants were presented with 60-s rhythmic sequences in both sensory modalities. The sequences were created in Matlab 2020b (MathWorks, MA, USA) by seamlessly looping 25 times a specific 2.4-s rhythmic pattern. The pattern was built by dividing 2.4 s into twelve equal intervals of 200 ms and arranging eight equivalent sensory events on this regular grid of time points, corresponding to [xxxx.xxx.. x.] (where x is a sensory event and dot an empty grid interval). The sensory events corresponded to 150-ms long sounds or vibrations (10-ms linear ramp-up, 90-ms plateau, and 50-ms linear ramp-down), followed by a 50-ms gap (see Figure 1A for a visualization of the sequential arrangement of sensory events composing this specific pattern).

**Figure 1.**
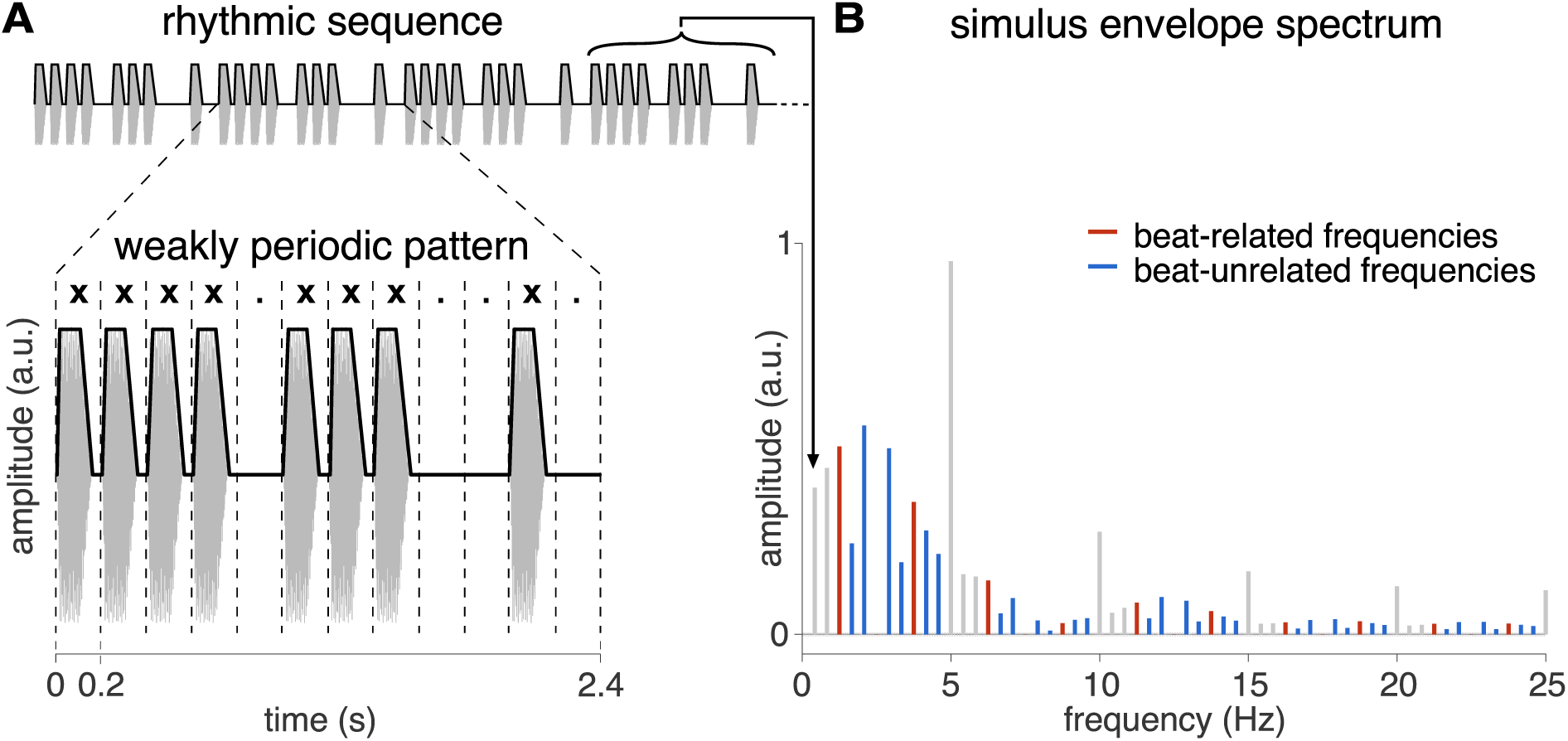
Stimuli consist in a repeated rhythmic pattern where the beat periodicity is not prominent. **A.** Time domain representation of the stimuli (full signal in grey, sound envelope in black). The 2.4-s rhythmic pattern was looped 25 times, yielding 60-s sequences for both sensory modalities (carrier frequency of 300 Hz and 86 Hz for acoustic and tactile sequences, respectively). **B.** Magnitude spectrum of stimulus envelope. Frequencies of interest were defined among the frequency of pattern repetition (1/2.4 s = 0.416 Hz) and harmonics. Harmonic frequencies corresponding to the most consistently tapped periods were defined as beat-related frequencies in red (4 x 200 ms = 800 ms i.e., 1.25 Hz and harmonics) and the remaining frequencies as beat-unrelated frequencies in blue. Note that beat-related frequencies (defined as the most consistently tapped period of 800 ms, i.e., 1.25 Hz and harmonics) were overall of lower magnitude than beat-unrelated frequencies in this rhythm.

This pattern, used in a number of previous studies (Nozaradan et al., 2016, 2017a, 2017b; Lenc et al. 2018; Sauvé et al., 2022; Sifuentes-Ortega et al., 2022), has been repeatedly shown to induce perception of a periodic pulse (beat) at a rate generally converging across Western participants towards a grouping of four underlying grid-intervals (4 x 200 ms). Moreover, significant relative enhancement of neural activity at frequencies corresponding to the rate of this beat periodicity and harmonics has been observed in the EEG in responses to this repeated pattern (Nozaradan et al., 2012; 2018; Lenc et al. 2022). Importantly, this specific pattern can be considered weakly periodic at the rate of this beat, since the groups of sensory events making up the rhythm are arranged in a way that does not prominently cue the beat periodicity (Poven and Essens 1985; Patel et al., 2005; Grahn et Brett 2007). Using a weakly-periodic rhythm is critical here, as it allows to control for low-level sensory confounds whereby the observed neural emphasis on the beat periodicity could otherwise be trivially explained by neural responses elicited by prominent physical features of the stimulus (see below and Figure 1B, for a quantification of the prominence of the beat periodicity in the stimulus modulation signal).

*Acoustic inputs.* Sound events consisted of pure sine waves at a carrier frequency of 300 Hz based on previous work having shown that stimuli within this frequency range elicit robust EEG responses (Evans and Deatherage 1969; Rahman et al., 2020; Nozaradan et al., 2015; 2018). Acoustic sequences were delivered binaurally at an intensity of 70 dB SPL using flat frequency response insert earphones (ER-2, Etymotic Research, IL, USA). *Tactile inputs*. Vibration events consisted of a 86-Hz sine waves at an intensity corresponding to a peak-to-peak displacement magnitude of 230 µm. Previous works have shown that such tactile stimuli effectively recruit the different types of mechanosensitive receptors and elicit robust EEG responses (Muniak et al. 2007; Bensmaïa et al., 2008; Rahman et al., 2020). Tactile sequences were generated by an electromagnetically shielded piezo-electric vibro-tactile stimulator (VTS, Arsalis, UCLouvain, Louvain-la-Neuve, Belgium) and delivered unilaterally to all fingertips in contact with a 20-mm diameter round-tipped probe. The specific cutaneous region of the fingertips was chosen given its highest density of mechanoreceptors (Johasson and Valbo 1979; Valbo and Johasson 1983; Corniani et al. 2020) and for its distance to the hearing canal which prevented any auditory response through bone or soft tissue conduction (Geal-Dor and Miriam 2021). Moreover, during tactile stimulation, the likelihood of eliciting auditory response to sounds produced by the piezo-electric stimulator was reduced by playing a uniformly distributed white noise through insert earphones at individually adjusted maximal tolerable intensity (up to 80 dB SPL). The masking white noise started 2 s before the onset of each tactile trial and ended 0.5 s after its end.

### Experimental design

The main experiment consisted of three blocks, each composed of an EEG session (twelve trials), followed by a tapping session (five trials). A short break of a few seconds was included between each trial and block to prevent sensory habituation and fatigue.

During the EEG session, participants were asked to focus their attention on the tactile or acoustic rhythmic sequences and refrain any movement. During tactile stimulation of the fingertips, position of the forearm and wrist was comfortably stabilized by means of a cushion. To further encourage participants to focus on the temporal properties of the stimuli, participants were also asked to detect transient changes of the tempo that could possibly occur in the stimulus sequence, and to report the presence and number of such changes at the end of each sequence. Tempo changes occurred in two non-consecutive trials pseudo-randomly placed within the twelve EEG trials composing each block (with the exclusion of the first trial, which never contained a tempo change). Tempo changes consisted of one pattern in which the underlying grid intervals were progressively lengthened (from 200 ms to 230 ms for the acoustic sequences, and from 200 ms to 250 ms for the tactile sequences), and then shortened back to the initial 200-ms grid intervals following a cosine function across the twelve grid intervals spanning one repetition of the pattern. Within the 60-s sequences containing the tempo change, up to three non-consecutive altered patterns were pseudo-randomly positioned among the 25 repetitions of the pattern, excluding the first repetition. The altered EEG trials were removed from further analyses.

During the tapping session, participants were asked to tap the perceived beat along with the rhythmic sequences over five successive trials, using the index finger of their preferred hand. Finger tapping was recorded using a custom-build response box (hereafter “tapping box”; Institute of Neurosciences, UCLouvain, Belgium) containing a high resistance switch able to generate a trigger signal every time the fingertip contacts the response box and a force sensor continuously monitoring the normal force applied to the box (with a constant response delay of 62.5 μs). The surface of the tapping box contacted by the finger was rigid, providing somatosensory and possibly auditory feedback. This feedback was reduced by using insert earphones, which partially blocked the sound of each tap during the presentation of the acoustic stimulus and completely masked it with white noise during tactile stimulation. Participants were asked to start tapping as soon as possible after the rhythmic sequence started and continuously tap the perceived beat along with the entire sequence as regularly as possible and as synchronized as possible with the sequence. Participants were advised not to restrict spontaneous movements of other body parts if doing so would help them perform the tapping task.

### Familiarization phase

The main experiment was preceded by a familiarization phase in which the participants were briefed on the concept of (i) beat, required for the tapping task, and (ii) tempo change, required for the detection task during the EEG recording sessions.

During the first part of the familiarization task, participants were asked to press the space bar of a keyboard with the index finger of their choice while listening successively to three tracks of electronic music in which the beat was acoustically either very explicit or more ambiguous. The instruction was to tap continuously along the track and as regularly as possible in synchrony with the beat they perceived, as they would do if they were nodding the head or stepping along with the music tracks. A plausible beat was indicated at the start of each music track by overlaid periodic hand-claps sounds that gradually faded out as the track progressed. Participants were asked to initially synchronize their taps to the clap sounds and keep tapping despite the cue was fading out (i.e., in a synchronization-continuation mode). Then, participants were presented with rhythmic sequences whose stimulus parameters were all identical to those used during the main experiment, except for the specific sequential arrangement of sensory events forming the repeated rhythmic pattern. Namely, we used a weakly periodic rhythmic pattern different from the one used in the main experiment, corresponding here to [xxx.xx..x.x.] (where x is a sensory event and dot an empty grid interval). These either acoustic or tactile 60-s familiarization-specific rhythmic sequences were presented while participants were encouraged to tap along the perceived beat along as regularly as possible on the tapping box, following the exact same instructions as in the main experiment. Tactile sequences were delivered to the hand contralateral to the hand chosen by the participants to perform the tapping task. Participants were explicitly asked to relax and keep their fingers still on the stimulator while the sequences were played.

In the second part of the familiarization phase, participants were accustomed with the detection of tempo changes as occurring in a few trials of each block in the main experiment. To this aim, participants were asked to detect tempo changes pseudo-randomly inserted in the familiarization-specific rhythmic sequences in the same fashion as in the main experiment.

### EEG recording and processing

The EEG was recorded using 64 sintered Ag-AgCl electrodes placed on the scalp according to the international 10/20 system (Active Two, Biosemi, The Netherlands). Two additional electrodes were placed on the left and right mastoids. The signal was referenced to the CMS (Common Mode Sense) electrode and digitized at a 1024-Hz sampling rate (with default hardware low-pass filtering at one fifth of the sampling rate). Electrode offsets were kept below 50 mV for all leads. The continuous EEG recordings were processed off-line with Letswave6 (https://www.letswave.org/) and custom scripts in Matlab 2020b (MathWorks, MA, USA). The continuous EEG signal was filtered using a 0.1-Hz Butterworth zero-phase high-pass filter (second order) to remove irrelevant slow fluctuations, and then segmented into 60-s epochs relative to trial onset, thus capturing the total duration of stimulation in each trial. Channels containing artifacts exceeding ± 200 mV or excessive noise were linearly interpolated using the three closest channels (a maximum of three channels were interpolated per participant in less than 7% of the total sample). After re-referencing to the common average, artifacts due to eyeblinks, eye movements, muscular activity or heartbeat were removed using independent component analysis (FastICA algorithm) (Hyvärinen and Oja 2000). A maximum of three independent components were removed by participant. EEG responses recorded during acoustic stimulation were analyzed at a fronto-central pool of electrodes (F1, FC1, C1, F2, FC2, C2, Fz, FCz, Cz) re-referenced to the averaged mastoids which is a standard reference to estimate cortical auditory responses (Skoe & Kraus, 2010; Mahajan, 2017; Nozaradan et al., 2016; 2018). EEG trials recorded during tactile stimuli were analyzed on a sensorimotor pool of electrodes contralateral to the stimulated hand (C2, C4, C6, T8, TP8, CP6, CP4, CP2, P2, P4, P6, P8) re-referenced to Fz electrode which is a standard reference to estimate cortical somatosensory responses (Tobimatsu, 1999; Cruccu, 2008; Moungou, 2016; Meinhold, 2022). EEG signals recorded during right hand stimulation were spatially flipped over midline as if all participants were stimulated on the left fingertips. For each participant and condition, time domain EEG signals were then averaged to enhance the signal-to-noise ratio of the neural response by attenuating the contribution of activities that were not time-locked to the stimulus (Mouraux et al. 2011; Nozaradan et al. 2011, 2012).

### Estimation of beat prominence in the brain responses: magnitude spectrum-based analysis

For each participant and condition, the averaged time domain EEG epochs were transformed in the frequency domain using a fast Fourier Transform (FFT), yielding frequency spectra ranging from 0 to 512 Hz with a frequency resolution of 0.0167 Hz (1/60 s). A local baseline correction was subtracted from each frequency bin in the resulting spectrum to minimize the contribution of local variations of noise inherent to EEG recordings. The baseline was defined as the average magnitude measured at −2 to −5 and +2 to +5 frequency bins relative to each frequency bin (Mouraux et al. 2011; Xu et al. 2017). Spectra were then averaged across all channels (average pool) and across modality-specific selections of channels (fronto-central pool for auditory and contralateral parietal pool for somatosensory).

#### Identification of the frequencies of interest

The frequencies at which the cortical responses were expected to be elicited were determined based on the repetition rate of the rhythmic pattern. Specifically, the frequencies of interest constituted the pattern repetition rate (0.416 Hz = 1/2.4 s) and harmonics, since this set of frequencies captures any response that is reliably and consistently elicited by the repeating rhythmic pattern. We used the temporal envelope of the 60-s sequence stimuli as computed with Hilbert transform (function ‘hilbert’ as implemented in Matlab 2020b). The resulting modulation signal was then transformed in the frequency domain using an FFT, yielding a frequency spectrum of the input envelope with a spectral resolution of 0.0167 Hz (figure 1B; Nozaradan 2017a, b). The range of included frequencies was adjusted for each modality, i.e., between 0 and 15 Hz for auditory responses and 0 and 25 Hz for somatosensory responses (see below for the determination of the response bandwidth for each modality). From the resulting set of harmonic frequencies, the first two frequencies were discarded from further analyses as they were located within a frequency range (< 1 Hz) typically featuring prominent 1/f-like background noise in EEG spectra, thus prone to unreliable measures (Cirelli et al., 2016; Lenc et al., 2023). Moreover, the twelfth frequency (i.e., 5 Hz) and its harmonics were also dismissed, as their magnitude is expected to be driven in major part by the shape of the individual 200-ms events composing the rhythmic pattern (i.e., 1/0.2 s = 5 Hz, and harmonics).

#### Estimation of the response frequency bandwidth for each modality

To determine the frequency range where the EEG responses for each modality were distributed, the magnitude obtained from the group-level averaged, baseline-corrected EEG spectra were summed successively over all harmonic frequencies from 0.416 Hz (i.e. 1/2.4 s) up to 30 Hz (72 frequencies) for each modality separately. All harmonic frequencies were taken into consideration, as the aim of this analysis was to quantify the overall response to the rhythmic input irrespective of any higher-level transformation. As shown in Figure 2, the curves of the summed magnitude as a function of frequency demonstrate a substantial gain in magnitude every 5 Hz. This prominent periodicity in the EEG signals reflects the responses to the recurring shortest inter-onset intervals making up the rhythmic pattern (200 ms inter-onset or grid intervals, i.e. 5 Hz rate). We then estimated the slope of each 5-Hz segment of the obtained curves by fitting a linear regression model. Finally, the response frequency bandwidth was determined for each modality by identifying the harmonic frequency at which the slope fell below an arbitrary threshold of 0.01 (i.e., a gain in magnitude less than 0.05 μV over a range of 5 Hz). The estimation of the response frequency bandwidth for the auditory modality was 0-15 Hz, with most of the response concentrated below 5 Hz, while the bandwidth of the somatosensory response was 0-25 Hz. This estimation was also confirmed by computing the derivative of the curve and selecting the frequency at which the derivative was minimal and tented towards zero. Only the frequencies of interest located within the obtained response frequency bandwidths were further used to compute the beat-related z-scores of the stimuli and the EEG responses for each modality respectively.

**Figure 2.**
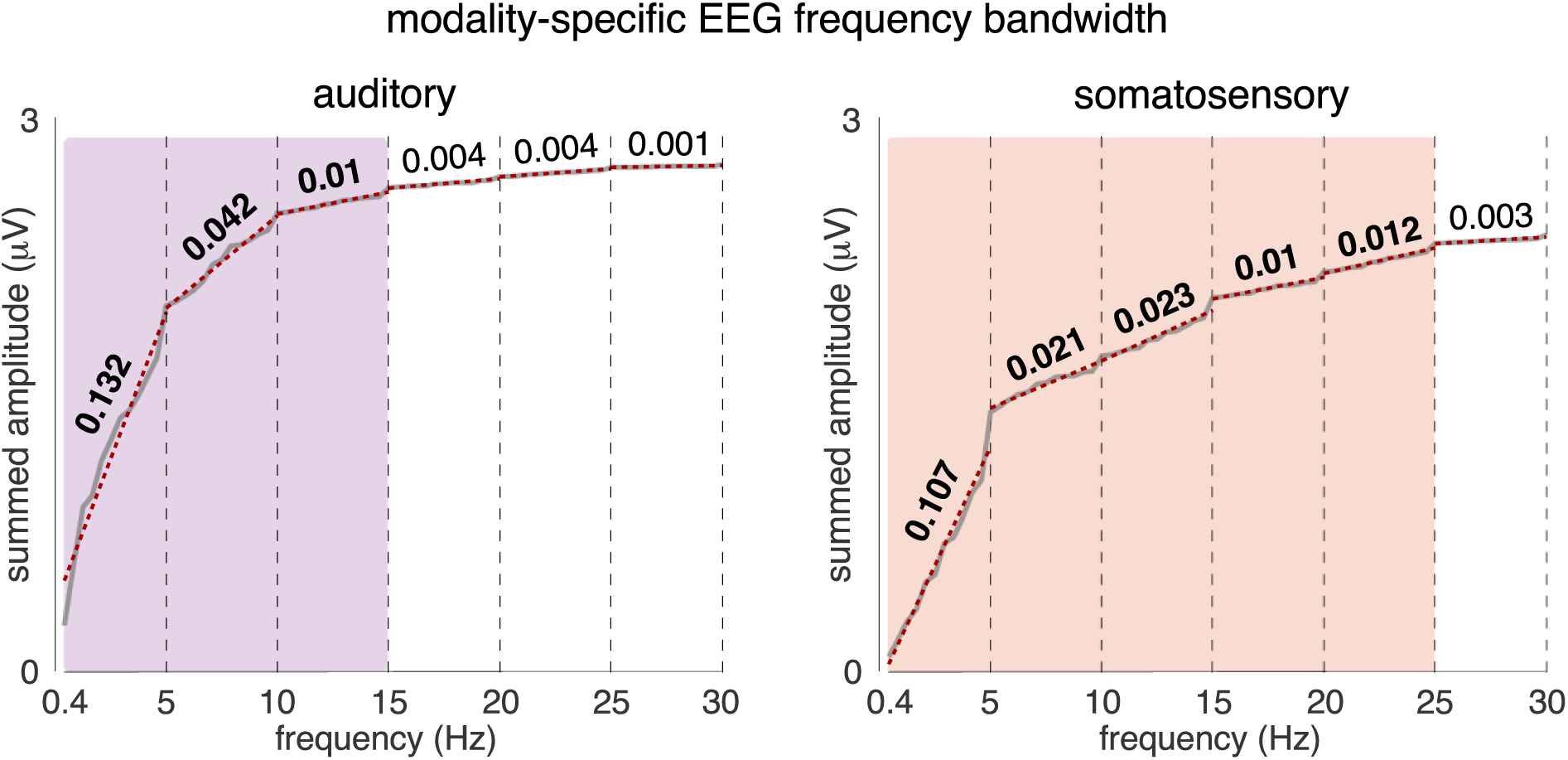
EEG responses show frequency bandwidths with lower cut-off for acoustic (0-15 Hz, in purple) than tactile (0-25 Hz, in green) input. The modality-specific bandwidths were determined based on the summed amplitudes over harmonic frequencies of pattern repetition rate, measured from the group-level averaged EEG magnitude spectrum, separately for each modality. Obtained summed amplitudes (grey curve) are superimposed red dashed lines corresponding to linear functions fitted to 5-Hz long segments across the spectra. The slopes of the fitted lines show faster convergence towards zero for the auditory EEG responses, revealing the lower cut-off of their bandwidth. Note that most of the auditory response is concentrated below 5 Hz.

#### Z-scored signal-to-noise ratio measurements (zSNR)

To ensure that cross-modal differences in magnitude at the frequencies of interest were not driven by different quality of the signal, z-scored signal to noise ratio (zSNR) was computed from the raw spectra (i.e., without baseline correction) to statistically test whether the response significantly stands out from background noise in the recorded signal (Liu-Shuang et al., 2014; Jonas et al., 2016; Lochy et al., 2018; Volfart et al., 2020; Hagen et al., 2021). First, the magnitude of at all frequency bins of interest and the respective local noise at the surrounding bins ranging from −12 to −2 and +2 to +12 were extracted. The zSNR value was then obtained as ([average magnitude across frequencies of interest] – [average noise magnitude]) / [standard deviation of the baseline magnitude].

#### Measurement of relative prominence of the beat-related frequencies

The main goal of the current study was to assess, from the whole set of frequencies of interest, the relative prominence of the frequencies considered as specifically related to the beat vs. the remaining frequencies which are included in the stimulus envelope magnitude spectrum but are unrelated to the beat periodicity. To this aim, the magnitude at each of these frequencies of interest was first standardized into z-scores (Lenc et al., 2020). The obtained z-scores were then averaged separately across beat-related frequencies (4 x 200 ms = 800 ms i.e., 1.25 Hz and harmonics) and beat-unrelated frequencies (i.e., the remaining frequencies of interest). An average beat-related z-score higher than the corresponding beat-related z-score calculated from the stimulus envelope spectrum (acoustic rhythm z-score = −0.059; tactile rhythm z-score = −0.051) would thus reflect selectively enhanced beat periodicity in the EEG compared to the input.

In addition, to assess whether the transformations observed in the EEG responses to the acoustic rhythm could not be trivially explained by responses at peripheral stages of sound processing, we also compared the recorded EEG responses to responses simulated with a biologically plausible model of peripheral auditory processing (see also Lenc et al., 2018; 2020). The cochlear model used to analyze the acoustic stimuli consisted in an Equivalent Rectangular Bandwidth (ERB) filter bank with 64 channels (Patterson & Holdsworth, 1996), followed by Meddis’ inner hair-cell model (Meddis, 1986), as implemented in the Auditory Toolbox for Matlab (Slaney, 1998). The output of the cochlear model was subsequently transformed into the frequency domain using FFT, and the obtained spectra were then averaged across cochlear channels. The beat-related z-score was then computed as described above (acoustic rhythm z-score from the model = −0.126, i.e. showing even less emphasis of the beat periodicities as compared to the acoustic signal itself).

### Estimation of beat prominence in the brain responses: autocorrelation-based analysis

A complementary approach was applied to analyze the prominence of beat-related periodicities in brain responses using autocorrelation (Lenc et al. 2024). This novel autocorrelation-based analysis aimed to corroborate the results obtained with the magnitude spectrum-based analysis described above. Critical to the current study, the autocorrelation-based approach has the advantage of providing an estimate of periodicity that is invariant to the shape of the recurring signal. This is of major importance here, because differences in the brain responses elicited by the acoustic vs. tactile sequences could be driven by differences unspecific to any actual beat-related periodization of the input but rather by lower-level properties of the response specific to the respective sensory modalities (e.g., cross-modality differences in the overall shape of the responses). As implemented here, the autocorrelation function (ACF) was estimated from the complex spectrum from which an estimate of the 1/f-like noise was estimated and subtracted. Then, only frequencies of interest were kept before computing the ACF. From this ACF, we extracted Pearson’s coefficients at lags of interest corresponding to (i) beat periodicities (beat-related lags: 0.8 s and multiples from which 2.4 s corresponding to the pattern duration was excluded), and (ii) control lags corresponding to periodicities where the beat was not perceived despite being compatible with the temporal arrangement of the sounds making up the rhythmic stimulus (beat-unrelated lags: 0.6 s, 1 s, 1.4 s and 1.8 s and multiples). After normalizing the coefficients across the whole set of lags using z-scoring, the periodicity of the response at the rate of the beat was quantified by averaging the coefficients across beat-related lags (for methodological details, see Lenc et al. 2024).

### Tapping recording and analysis

The tap onsets generated by the tapping box were sent as analog triggers to the EEG system and recorded at the same sampling rate as the EEG signal (i.e., 1024 Hz). The force signal continuously monitored by the tapping box was digitalized at 44100 Hz and recorded by means of an audio interface (Fireface UC, RME, Germany). The time series of the continuous force signal were down-sampled to 1024 Hz after having been low pass filtered at 300 Hz (i.e., below the Nyquist frequency of the target sampling rate) to avoid aliasing. Time series of tap onsets were generated as continuous time-domain signal with duration corresponding to the length of the stimulus sequence and sampled at 300 Hz. The value of each sample corresponding to a tap onset time was set to 1 (i.e., a unit impulse) and 0 otherwise.

#### Circular analysis of tapping

To estimate the beat periods most consistently tapped across participants, the median intertap interval (ITI) was computed using the tap onset times (i.e., times at which the finger contacted the tapping box) for each trial, block, and participant separately. Because participants typically waited a few sensory events before starting to tap along with the rhythmic input, the first 2.4-s (i.e., temporal window of the first pattern presentation) was discarded.

To quantify the tapping performance of each participant, the beat tapping stability was evaluated by calculating a circular measure, namely the mean vector length (Nozaradan et al. 2016, Berens 2009). This measure was calculated by first selecting, based on the median ITI of each participant, the closest plausible beat period, i.e., the beat period most likely targeted by the participant (with plausible beat periods corresponding here to any integer multiple of the 200-ms grid interval that would fit within the 2.4-s rhythmic pattern, thus yielding six plausible beat periods in total: 200 ms, 400 ms, 600 ms, 800 ms, 1200 ms or 2400 ms). The obtained target beat period was then used to compute a time series of target beat positions, with phase zero set in accordance with the first tap of each trial. The signed differences between each tap and the closest target beat position was then converted into an angle and mapped onto a unit circle. The resulting unit vectors were then average across trials and the length of the mean vector served as an index of beat stability. This mean vector length thus reflected the *consistency* of the asynchrony between taps and the corresponding beat positions, i.e., the strength of frequency locking between the beat and the tapping response (Rosenblum et al. 2001). To compare the tapping stability between blocks and participants, the angle of the mean vector was subtracted from the unit vectors corresponding to individual taps separately for each trial, the data were collapsed across trials, and the mean vector was recalculated to obtain one stability value per block for each participant.

#### Magnitude spectrum-based analysis of tap onsets and tap force

As a complementary estimate of tapping stability, single trials of (i) time series of tap onsets and (ii) continuous force signals as recorded by the tapping box were also transformed in the frequency domain using FFT. For each block and participant, and similarly to the analysis described above for stimulus input and EEG responses, z-scores of beat-related and beat-unrelated frequencies were then calculated from the magnitude spectrum obtained for each trial and averaged across trials. The frequency ranges of interest were defined as for EEG responses using the changes in slopes of the successive 5-Hz chunks obtained by successively summing the magnitude across harmonics of the pattern duration as computed on the group-level averaged magnitude spectrum for each type of signal (time series of tap onsets and tap force) and modality. Tapping responses were mainly distributed between 0 and 10 Hz for both type of signal and both sensory modalities.

#### Autocorrelation-based analysis of tap onsets and tap force

The relative prominence of beat periodicities was also assessed in time series of tap onsets and tap force using the novel implementation of the frequency-tagging approach based on autocorrelation as described above for EEG responses (Lenc et al. 2024).

### Head movements control

To rule out artefacts related to unintentional periodic head movements of participants synchronized to the perceived beat during the EEG recording, which may potentially enhanced beat-related periodicities in the obtained EEG responses, head movements were monitored by means of a 2-axes accelerometer (x for left-right, y for back-front) strapped on the EEG cap at the vertex. Signals were acquired at a 1024-Hz sampling rate. The relative prominence of beat-related frequencies was estimated in the accelerometer data in the same way as for tapping data.

### Statistical analyses

The statistical analyses were conducted using JASP (JASP Team 2023, Version 0.17.2). First, we verified that the data followed a normal distribution using Shapiro-Wilk tests. To assess if the order of blocks influenced the different measures obtained from EEG or tapping, mixed repeated measures analyses of variance (RM-ANOVAs) were performed with “Group” as a between-subject factor (group 1: acoustic-tactile-tactile vs. group 2: tactile-acoustic-tactile vs. group 3: tactile-tactile-acoustic) and “Block” as a within-subject factor (acoustic block vs. first tactile block vs. second tactile block). For between subject comparisons, homogeneity of variance was verified using Levene’s test. Sphericity was tested using Mauchly’s test and F-values were Greenhouse–Geisser corrected when this assumption was violated. Post-hoc comparisons were then performed using t-tests, with Holm correction for multiple comparisons. In case of non-parametric testing, Friedman tests were performed with the factor “Block”, with Conover’s tests used for post hoc comparisons. In addition to this frequentist analysis, we also calculated Bayes factors (BF_10_) to quantify the probability of the data under different models (Rouder et al. 2017, van den Bergh et al. 2020). Then, the inclusion Bayes factor (BF_incl_) was estimated for each model’s predictor to quantify the evidence in favor of including each of those predictors in the best model.

Finally, to assess the periodization of the EEG responses with respect to the input, one sample t-tests were performed between the beat-related z-scores of the EEG and the beat-related z-scores of the stimuli for each modality.

### Code Accessibility

Code for the autocorrelation-based approach analysis is available in OSF at https://osf.io/preprints/psyarxiv/yptsr. Acoustic and tactile stimuli are available upon request.

## Results

### Tapping data

#### Convergent beat periodicities across modalities revealed by tapping

Median intertap intervals (ITI) pointed toward convergent beat periodicities at 800 ms (i.e. corresponding to the grouping of four grid intervals, or 3 beats spanning one pattern repetition). Subject-level median ITIs were not significantly different between groups corresponding to different orders of block presentation (main effect of group: F_(2,42)_ = 0.454; p = 0.638; η_p_^2^ = 0.021; RM-ANOVA). Moreover, the median ITIs did not differ between blocks (Figure 3A; main effect of block: F_(1.3,54.73)_ = 2.412; p = 0.118; η_p_^2^ = 0.054), and no significant Group × Block interaction was observed (F_(2.6,54.73)_ = 0.573; p = 0.611; η_p_^2^ = 0.027). These results were corroborated by a Bayesian RM-ANOVA which showed that the null model better explained the data than any other model including the Group and Block factors and their interaction (all BF_10_ ≤ 0.584).

**Figure 3.**
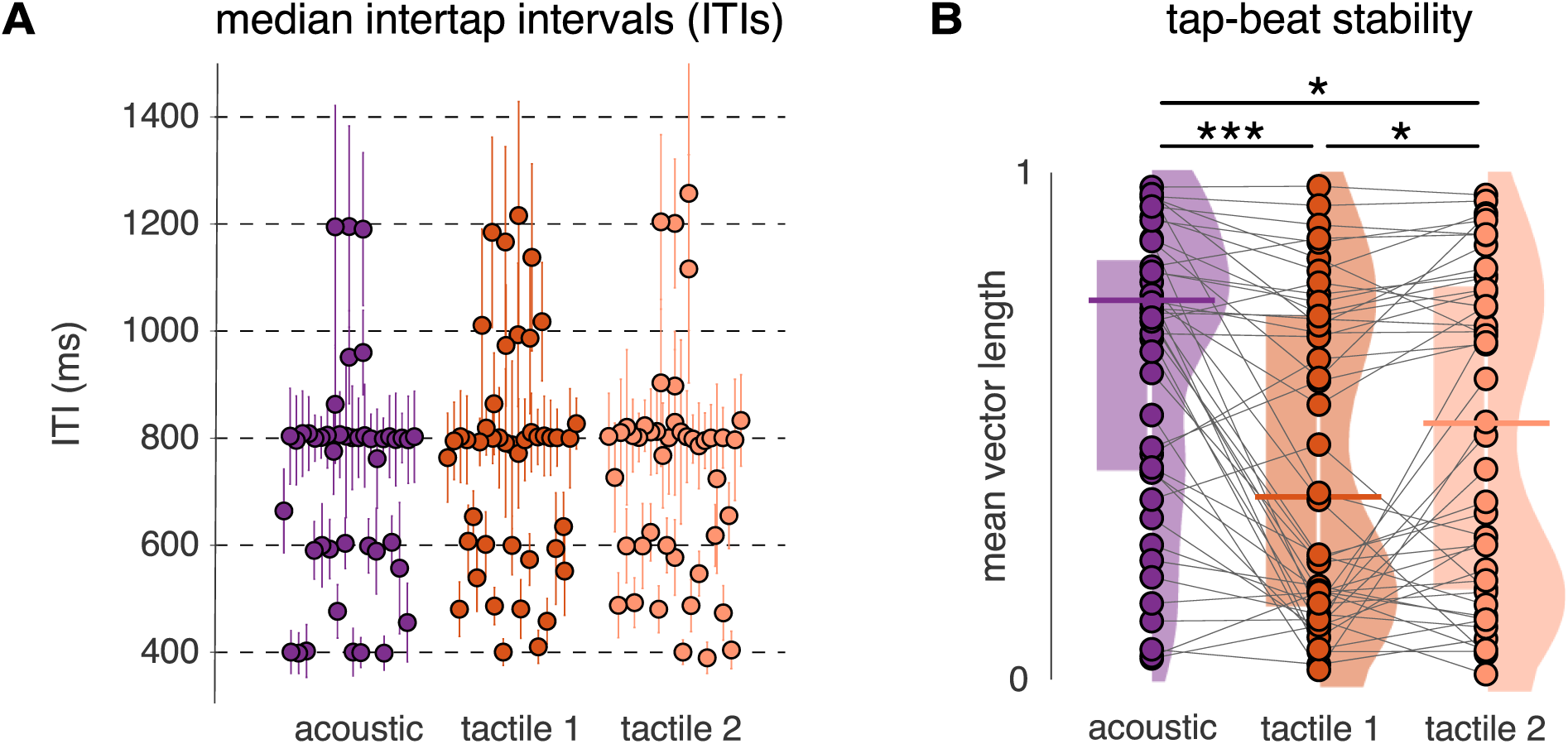
The tapping task reveals significantly lower beat tapping stability in response to the tactile vs. acoustic rhythm. **A.** Subject-level median intertap intervals (ITIs; one dot per participant; error bars indicate interquartile ranges) for each block condition. The tapped beat periods converged towards 800 ms across all blocks (4 x 200-ms grid interval, i.e., 3 beats per pattern repetition). One data point around 2.4 s in the three blocks is omitted from the plot for visualization purposes. **B.** Tapping stability (one dot per participant; horizontal bars indicate the median; left shaded boxes indicate interquartile ranges; fitted distribution densities are displayed on the right, separately for each block). Tapping stability was significantly reduced in the tactile conditions as compared to the acoustic condition (i.e., resultant vector length of the circular distribution of asynchronies between taps and beat positions closer to zero). Note that both panels A and B depict each block, irrespective of the order in which they were presented (given the absence of order effect). Asterisks indicate significant differences between blocks obtained from the Conover’s post-hoc comparisons (*p < 0.05, ***p < 0.001).

#### Reduced beat periodicities in tapping to tactile vs. acoustic inputs

*Beat tapping stability.* The obtained beat tapping stability deviated from a normal distribution (Shapiro-Wilk, p-values < 0.002), which justified further use of Friedman tests for comparison across blocks and groups, and Conover’s tests for post hoc comparisons. The Friedman test revealed a significant effect of Group (corresponding to different orders of block presentation; χ^2^_(2)_ = 18.711; p = 8.648×10^−5^; W = 0.208). Conover’s post-hoc comparisons showed significant differences between blocks, namely with larger stability for the acoustic block as compared to the first tactile (p = 1.218×10^−4^) and second tactile block (p = 0.048) and between the two tactile blocks (p = 0.045). These results were corroborated by a Bayesian RM-ANOVA which showed that the model including the Block factor (evidence for Block effect BF_incl_ = 7403.78) better explained the data than the null model or any models including the Group factor (all BF_10_ ≤ 0.366).

*Beat prominence in tap onsets time series.* The mean z-scored magnitudes at beat-related frequencies obtained using magnitude spectrum-based analysis were not affected by the group order of the blocks (F_(2,42)_ = 0.168; p = 0.846; η_p_^2^ = 0.008; RM-ANOVA). There was a significant effect of block (F_(1.57,65.95)_ = 12.405; p = 1.057×10^−4^; η_p_^2^ = 0.228; RM-ANOVA), and no significant Group × Block interaction (F_(3.14,65.95)_ = 0.360; p = 0.791; η_p_^2^ = 0.017). Post-hoc comparisons showed that beat-related frequencies were significantly more prominent in the acoustic block as compared to the first (t_(44)_ = 4.733; p = 2.663×10^−5^; Cohen’s d = 0.450) and second tactile blocks (t_(44)_ = 3.712; p = 7.386×10^−4^; Cohen’s d = .353; paired t-test), and the two tactile blocks were not significantly different from each other (t_(44)_ = −1.021; p = 0.310; Cohen’s d = −0.097; paired t-test; see Figure 4A). These results were corroborated by a Bayesian RM-ANOVA which showed that the best model included only the Block factor (effect of Block BF_incl_ = 1397.7) and better explained the data than the null model or any models including the Group factor (all BF_10_ ≤ 0.379).

**Figure 4.**
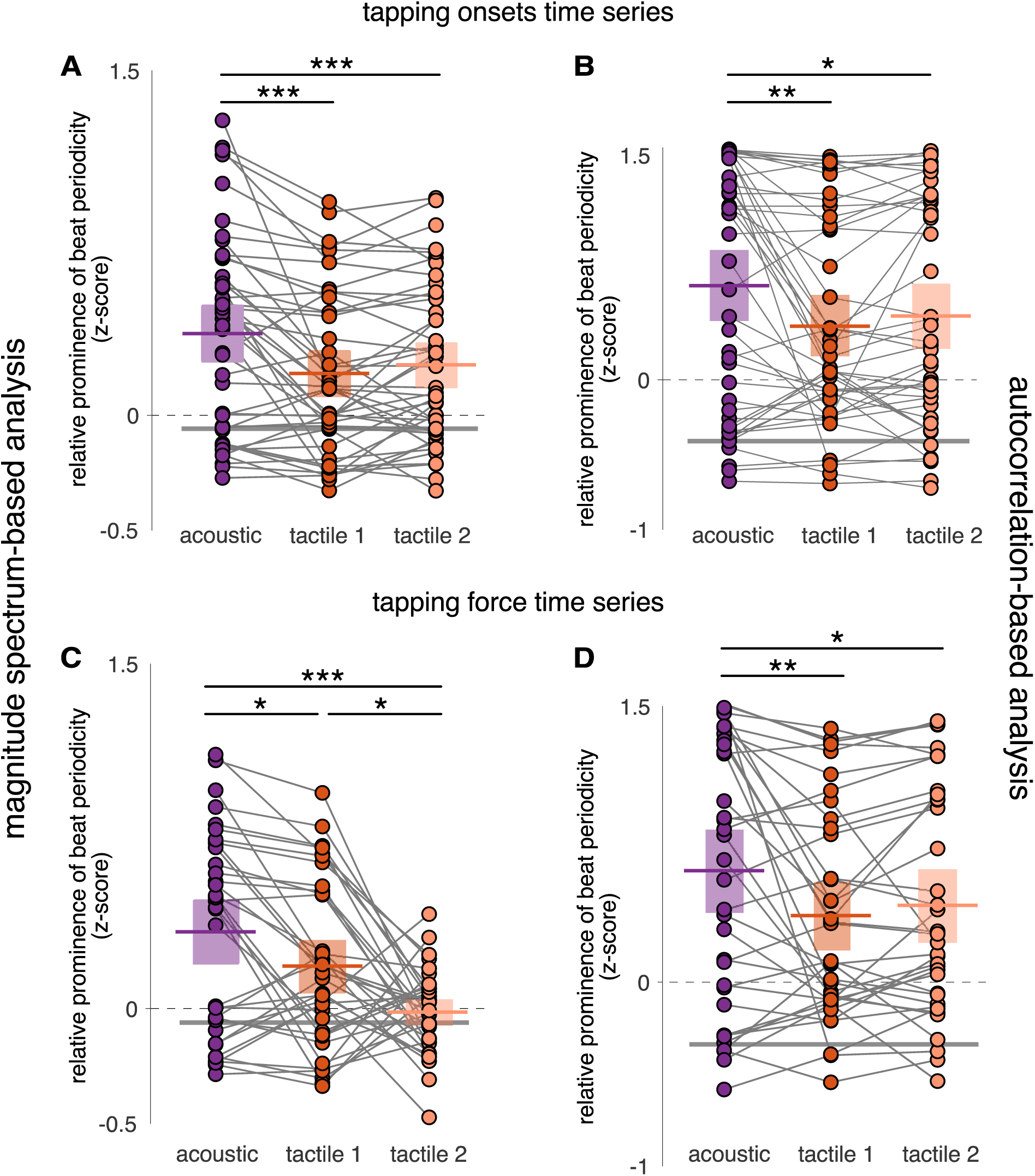
Tapping data show greater prominence of the beat periodicity in response to acoustic vs. tactile rhythm. Tap onsets and tapping force are depicted in A - B and C - D, respectively. Magnitude spectrum-based and autocorrelation-based analyses are depicted in A - C, and B - D, respectively. Note the significantly greater periodization of acoustic vs. tactile input (one dot per participant; colored horizontal bars indicate means; boxes show 95% confidence intervals; grey bars for corresponding values from stimulus envelope). Asterisks indicate significant differences between blocks obtained from the pairwise post-hoc comparisons (*p < 0.05, **p < 0.01, ***p < 0.001).

Those results were also confirmed using beat-related z-scored values obtained by the autocorrelation-based analysis at beat-related lags (Figure 4B). There was no significant main effect of Group (F_(2,42)_ = 1.0; p = 0.376; η_p_^2^ = 0.045; RM-ANOVA), a significant effect of Block (F_(1.49,62.43)_ = 7.487; p = 0.003; η_p_^2^ = 0.151; RM-ANOVA), and no significant Group × Block interaction (F_(2.97,62.43)_ = 0.572; p = 0.634; η_p_^2^ = 0.027). Post-hoc comparisons showed that beat-related lags were significantly more prominent in the acoustic block as compared to the first (t_(44)_ = 3.721; p = 0.001; Cohen’s d = 0.360) and second tactile blocks (t_(44)_ = 2.781; p = 0.013; Cohen’s d = 0.269; paired t-test), and the two tactile blocks were not significantly different from each other (t_(44)_ = −0.940; p = 0.350; Cohen’s d = −0.091; paired t-test). These results were corroborated by a Bayesian RM-ANOVA which showed that the best model included only the Block factor (effect of Block BF_incl_ = 28.46) and better explained the data than the null model or any models including the Group factor (all BF_10_ ≤ 0.551).

*Beat prominence in force signal time series.* Due to a technical problem, the force signal was recorded in 36 participants out of 45. Nevertheless, the analyses of beat prominence using the force signal converged with those performed on tap onsets. Mean z-scores obtained from the magnitude spectrum-based analysis were not affected by the group order of the blocks (F_(2,33)_ = 0.335; p = 0.718; η_p_^2^ = 0.020; RM-ANOVA). There was a significant effect of Block (F_(1.68,55.43)_ = 11.768; p = 1.360×10^−4^; η_p_^2^ = 0.263; RM-ANOVA), and no significant Group × Block interaction (F_(3.36,55.43)_ = 2.106; p = 0.103; η_p_^2^ = 0.113). Post-hoc comparisons showed that beat-related frequencies were significantly more prominent in the acoustic block as compared to the first tactile (t_(44)_ = 2.070; p = 0.042; Cohen’s d = 0.399; paired t-test) and second tactile blocks (t_(44)_ = 4.835; p = 2.495×10^−5^; Cohen’s d = 0.931; paired t-test). As compared to the analyses of onsets times series, the only noticeable difference was the significant decrease of beat-related z-scores in the second vs. first tactile block (t_(44)_ = 2.765; p = 0.015; Cohen’s d = 0.532; paired t-test; see Figure 4C). The effect of block was supported by a Bayesian RM-ANOVA which showed that the best model included only the Block factor (effect of Block BF_incl_ = 3680.16) and better explained the data than the null model or any models including the Group factor (all BF_10_ ≤ 0.216).

The results on force time series were confirmed using the mean z-scored values obtained with the autocorrelation-based analysis at beat-related lags (Figure 4D), also in line with the results obtained from the autocorrelation-based analysis of tap onsets (Figure 4B). There was no significant main effect of Group (F_(2,33)_ = 1.625; p = 0.212; η_p_^2^ = 0.073; RM-ANOVA), a significant effect of Block (F_(1.42,46.91)_ = 6.161; p = 0.009; η_p_^2^ = 0.157; RM-ANOVA), and no significant Group × Block interaction (F_(2.84,46.91)_ = 0.27; p = 0.836; η_p_^2^ = 0.016). Post-hoc comparisons showed that beat-related lags were significantly more prominent in the acoustic block as compared to the first (t_(44)_ = 3.362; p = 0.004; Cohen’s d = 0.412) and second tactile blocks (t_(44)_ = 2.555; p = 0.026; Cohen’s d = 0.313; paired t-test), and the two tactile blocks were not significantly different from each other (t_(44)_ = −0.807; p = 0.422; Cohen’s d = −0.099; paired t-test). These results were corroborated by a Bayesian RM-ANOVA which showed that the best model included only the Block factor (effect of Block BF_incl_ = 7.186) and better explained the data than the null model or any models including the Group factor (all BF_10_ ≤ 0.750).

### EEG responses

#### Reduced beat periodicities in brain responses to tactile vs. acoustic rhythms

The mean z-scored beat-related frequencies obtained using the magnitude spectrum-based approach were not affected by the group order of the blocks (F_(2,42)_ = 0.661; p = 0.522; η_p_^2^ = 0.031; RM-ANOVA). There was a significant effect of the Block (F_(1.94,81.46)_ = 37.044; p = 5.85×10^−12^; η_p_^2^ = 0.469; RM-ANOVA) and a significant Group × Block interaction (F_(3.88,81.46)_ = 3.769; p = 0.008; η_p_^2^ = 0.152). Post-hoc comparisons showed that beat-related frequencies were significantly more prominent in the acoustic block as compared to the first (t_(44)_ = 8.240; p = 6.008×10^−12^; Cohen’s d = 1.648; paired t-test) and second tactile blocks (t_(44)_ = 6.274; p = 2.927×10^−8^; Cohen’s d = 1.255; paired t-test) which were not significantly different from each other (t_(44)_ = −1.966; p = 0.158; Cohen’s d = −0.393; paired t-test; see Figure 6A). These results were supported by a Bayesian RM-ANOVA which showed that the model that better explained the data included both Block and Group factors and their interaction as compared to any other models (BF_10_ ≤ 0.575). There is decisive evidence for the effect of Block (BF_incl_ = 1.956×10^10^) followed by strong evidence for the Block × Group interaction (BF_incl_ = 10.991) and no evidence for including the Group factor (BF_incl_ = 0.158).

#### Periodization of the acoustic but not tactile input in the EEG

To assess the periodization of the input in the EEG responses, one-sample t-tests were performed between the beat-related z-scores of the EEG responses and the corresponding stimulus, separately for each modality. For the auditory modality, z-scored magnitudes of beat-related frequencies in the EEG response were significantly larger than in the stimulus beat-related z-score calculated from the envelope spectrum (t_(44)_ = 6.907; p = 7.818×10^−9^; Cohen’s d = 1.030; one-sided one-sample t-test against stimulus z-score of −0.059) and from the cochlear model (t_(44)_ = 9.567; p = 1.283×10^−12^; Cohen’s d = 1.426; one-sided one-sample t-test against stimulus z-score of −0.126). This was not the case in the somatosensory modality (tactile 1 block: t_(44)_ = −5.258; p = 1.00; Cohen’s d = −0.784; and tactile 2 block: t_(44)_ = −0.815; p = 0.790; Cohen’s d = −0.121; one-sided one-sample t-tests against stimulus z-score of −0.051).

#### Autocorrelation based analysis corroborated the magnitude spectrum analysis showing reduced beat periodicities in brain responses to tactile vs. acoustic rhythms

Importantly, the results of beat prominence in the EEG as obtained with the magnitude spectrum-based analysis were confirmed by the autocorrelation-based analysis (Figure 6B). The mean z-scored magnitudes at beat-related lags were not affected by the group order of the blocks (F_(2,42)_ = 2.263; p = 0.117; η_p_^2^ = 0.097; RM-ANOVA). There was a significant effect of Block (F_(1.81,76.05)_ = 12.464; p = 3.879×10^−5^; η_p_^2^ = 0.229; RM-ANOVA) and a significant Group × Block interaction (F_(3.62,76.05)_ = 2.121; p = 0.093; η_p_^2^ = 0.092). Post-hoc comparisons showed that beat-related lags were significantly more prominent in the acoustic block as compared to the first tactile (t_(44)_ = 4.626; p = 4.02×10^−5^; Cohen’s d = 0.960; paired t-test) and second tactile (t_(44)_ = 3.940; p = 3.362×10^−4^; Cohen’s d = 0.818; paired t-test) blocks which were close to be significantly different from each other (t_(44)_ = −0.687; p = 0.494; Cohen’s d = −0.143; paired t-test). These results were supported by a Bayesian RM-ANOVA which showed that the model that better explained the data included the Block factor as compared to any other models (BF_10_ ≤ 0.41). Indeed, there was decisive evidence for the effect of block (BF_incl_ = 3181.42), weak evidence for the Block × Group interaction (BF_incl_ = 1.091), and no evidence for including the Group factor (BF_incl_ = 0.376).

Similarly, as in the magnitude spectrum-based analysis, the beat-related z-scores obtained using the autocorrelation-based analysis showed a lack of periodization of the input in the EEG response to the tactile rhythm as opposed to the acoustic rhythm. Z-scored magnitudes of beat-related frequencies in the EEG response to the acoustic rhythm were significantly larger than in the stimulus z-scores obtained from the envelope spectrum (t_(44)_ = 6.495; p = 3.156×10^−8^; Cohen’s d = 0.968; one-sided one-sample t-test against stimulus z-score of −0.409) and from the cochlear model (t_(44)_ = 6.76; p = 1.288×10^−8^; Cohen’s d = 1.008; one-sided one-sample t-test against stimulus z-score of −0.432). This was not the case for EEG responses to the tactile rhythm (tactile 1 block: t_(44)_ = −0.004; p = 0.502; Cohen’s d = −6.544×10^−4^, and tactile 2 block: t_(44)_ = 0.848; p = 0.201; Cohen’s d = 0.126; one-sided one-sample t-tests against stimulus z-score of −0.409).

### Additional control analyses

To control for potential biases in our analyses, we conducted several additional analyses. *Ruling out confounds with overall magnitude of the EEG responses.* To ensure that any differences observed between conditions were not trivially explained by differences in the overall magnitude of the responses irrespective of any beat-related periodization of the input, z-scored signal-to-noise ratio measurements (zSNR) were computed for each block by pooling over magnitudes at all frequencies of interest (i.e. including all frequencies tagged as beat-related and beat-unrelated, within the response frequency bandwidth specific to each modality). The obtained zSNR values significantly deviated from a normal distribution (Shapiro-Wilk, p-values ≤ 0.029). The Friedman test revealed a significant effect of block (χ^2^_(2)_ = 10.711; p = 0.005; W = 0.119). Conover’s post-hoc comparisons showed significant differences between the acoustic and first tactile blocks (p = 0.005) and no significant difference between acoustic and second tactile blocks nor between tactile blocks (p-values = 0.153). Therefore, the differences in overall signal magnitude could not directly explain the cross-modal differences in beat-related periodization observed in the EEG data.

*Excluding contribution of unintentional head movement artifacts in EEG responses*. To rule out the possibility that beat-related periodization of the input in the EEG was driven by artefacts related to unintentional head movements of participants during the EEG recording, we estimated the prominence of beat periodicities in the accelerometer data (averaged across the two accelerometer axes) during EEG recording using the magnitude spectrum-based analysis. Due to a technical problem, accelerometer data were available for 41 participants out of 45. We compared the obtained z-scored magnitude of beat periodicities in the head movements to the corresponding values from the stimulus. There was a significant increase of beat periodicities in each block: acoustic (t_(40)_ = 2.704; p = 0.010; Cohen’s d = 0.420; one-sided one-sample t-test against stimulus z-score of −0.126), first tactile (t_(40)_ = 3.306; p = 0.002; Cohen’s d = 0.516; one-sided one-sample t-test against stimulus z-score of −0.051) and second tactile (t_(40)_ = 2.865; p = 0.007; Cohen’s d = 0.447; one-sided one-sample t-test against stimulus z-score of −0.051). However, there was no significant main effect of Block (F_(1.65,62.69)_ = 0.255; p = 0.733; η_p_^2^ = 0.007; RM-ANOVA), Group (F_(2,38)_ = 1.045; p = 0.362; η_p_^2^ = 0.052; RM-ANOVA), or Block × Group interaction (F_(3.3,62.69)_ = 1.741; p = 0.163; η_p_^2^ = 0.084; RM-ANOVA). Therefore, the unintentional head movements produced during EEG recordings were unlikely to explain the cross-modal differences in beat-related periodization observed in the EEG.

*Modality specific vs. common average montages for EEG analyses*. To ensure that the choices of the modality-specific pools of EEG channels and corresponding re-referencing did not bias the results, the same magnitude spectrum-based analysis of beat prominence in the EEG responses was performed on the signal extracted from all EEG channels re-referenced to the common average for both modalities. The results converged with the analyses performed on signals obtained from modality-specific montages. There was a significant main effect of Block (F_(1.79,75.29)_ = 38.586; p = 1.299×10^−12^; η_p_^2^ = 0.479; RM-ANOVA), no significant main effect of Group (F_(2,42)_ = 0.082; p = 0.921; η_p_^2^ = 0.004; RM-ANOVA), and no significant Block × Group interaction (F_(3.58,75.29)_ = 0.198; p = 0.924; η_p_^2^ = 0.009; RM-ANOVA).

## Discussion

The current study shows that mapping periodic beats onto an acoustic rhythm is related to enhanced beat periodicities in the neural representation of the rhythm. Moreover, this neural enhancement of the beat selectively projects onto a low-frequency range (under 15 Hz, mainly under 5 Hz). In contrast, presenting the same rhythm through the somatosensory modality does not produce such a periodic neural emphasis, despite significant and comparably robust neural responses to the tactile rhythm. Importantly, this cross-sensory difference converges with differences in the ability to tap the beat along with the acoustic vs. tactile rhythm.

In sum, internal representations of rhythm which might be experienced as *the beat* seems preferentially supported by periodized low-frequency neural activity. However, these higher-level neural representations are not necessarily shared across the senses, as well as the ability to move to the beat. Such periodized low-frequency neural activity may thus reflect temporal integration across multiple timescales beyond onset timing, a distinctive specialization of the auditory system over other sensory modalities, that supports higher-level internal representation and motor coordination with rhythm.

### Periodized neural and behavioral representation of acoustic vs. tactile rhythm

Most studies that investigated rhythm perception and sensorimotor synchronization across the senses have focused on instances where synchronization was meant to be performed in a one-to-one manner with the rhythmic input (Gilmore and Russo 2021, Ammirante et al. 2016, Tranchant et al. 2017). In other words, the goal was to synchronize each movement with the onset of each sensory event. In contrast, moving to the perceived beat along with the rhythms used here, where the beat is not prominently cued, implies to go beyond such one-to-one mapping. Namely, it requires an internal representation showing higher degree of invariance with respect to the temporal structure of the rhythmic input. Notably, this phenomenon is not a peculiarity of experimental design constrains or some specific music genres, but which abounds in music and dance worldwide (Butler, 2006; London, 2012; Margulis, 2014; London et al., 2017; Witek, 2017; Câmara and Danielsen, 2018). In the current study, the neural responses to the acoustic but not tactile rhythm show a specific, behavior-relevant, periodized representation of the input. This result thus adds to the growing evidence showing that such a selective neural emphasis could reflect internal templates of periodic beats beyond mere lower-level sensory confounds (Nozaradan et al., 2017, Lenc et al., 2021, Tal et al., 2017).

Another important observation is that a vast majority of participants spontaneously tap the beat at convergent periods, whether the rhythm is acoustic or tactile. However, while the tapping period is shared across sensory modalities, beat stability and prominence in the tapping are significantly lower for tactile vs. acoustic rhythm. Both the lower stability and beat prominence in the tapping, combined with the lack of emphasis of the beat in the EEG activity, thus suggests a functional link between the two measures. Yet, neural and behavioral observations offer a window onto different processes recruited over very contrastive tasks – namely, experiencing a rhythm while being instructed to stay still vs. actively producing synchronized movements – which might differently affect the nature of the underlying internal representations emerging during each of these tasks, respectively (Su and Pöppel, 2012; Manning and Schutz 2013). Nonetheless, while plausibly linked, these measures do not capture the underlying processes in a strict one-to-one fashion, thus highlighting their complementarity in understanding beat perception.

### Low-frequency neural activity supports periodization of acoustic but not tactile rhythm

In the current study, neural response to the acoustic rhythm selectively projects onto the low-frequency range, mainly under 5 Hz. In the time domain, this low-frequency activity manifests as slow fluctuations punctuated by more transient responses to each event onset (Fig. 5). This low-frequency activity could be a feature enabling the auditory system to integrate fast incoming events into slower, behavior-relevant, temporal units, which is critical to further coordinate body movement with rhythmic inputs. Importantly, this observation is in line with the crucial role of delta (< 4 Hz) and theta (4-8 Hz) bands oscillations in subserving multi-scale temporal integration of temporally structured input such as music and speech in humans (Arnal et al., 2015; Doelling et al. 2015; Teng et al. 2016; 2018) and non-human primates (Lakatos et al., 2005; 2016; Schroeder and Lakatos 2009).

**Figure 5.**
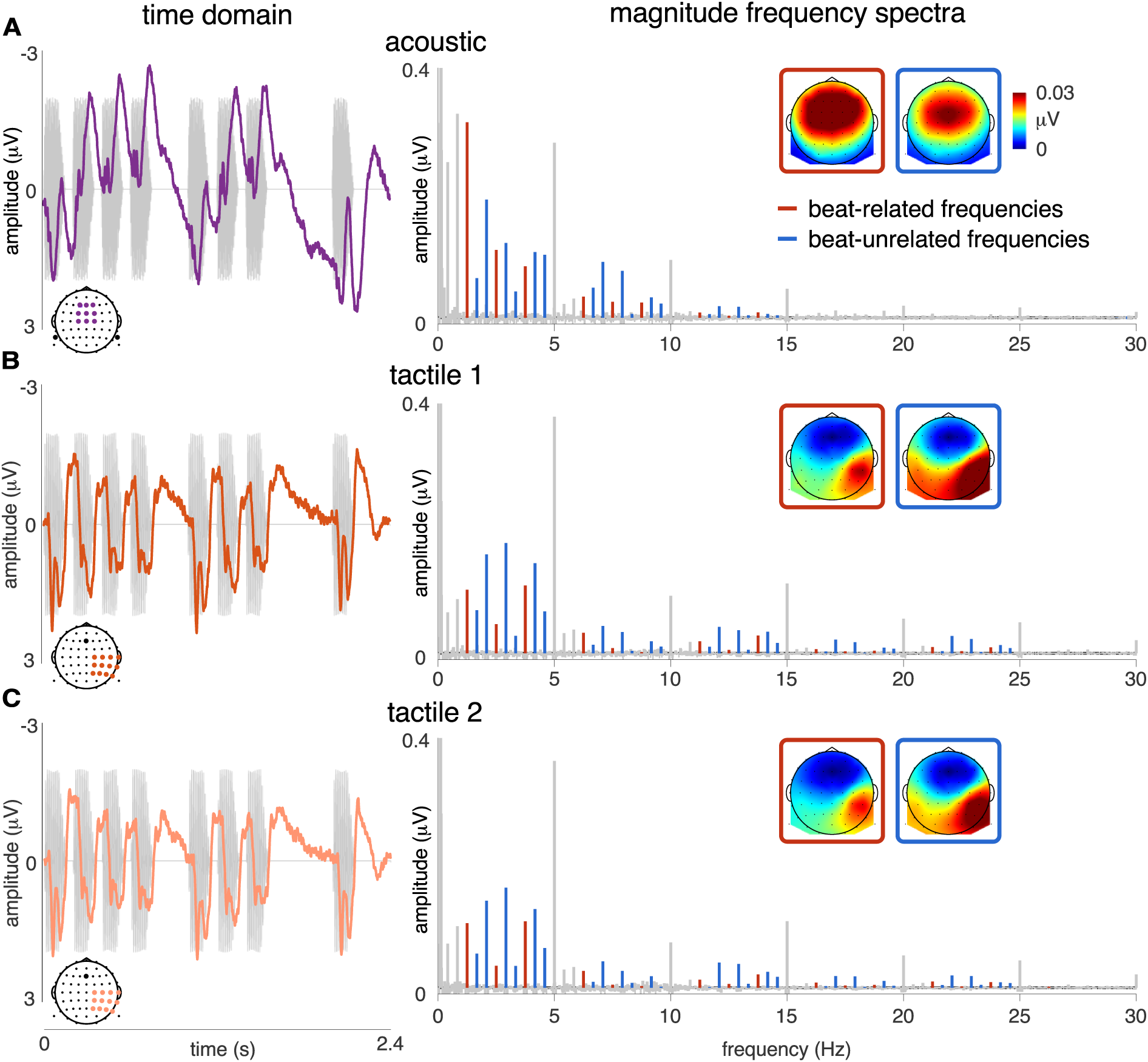
EEG responses in the time (left) and frequency domains (right) show slower fluctuations (lower-frequency) in response to the acoustic rhythm and more transient (higher-frequency) responses to tactile rhythm. Grand-average EEG responses in the acoustic (row **A**; fronto-central pool of electrodes referenced to the averaged mastoids), and first and second tactile blocks (rows **B** and **C**, respectively; sensorimotor pool of electrodes contralateral to the stimulated hand referenced to Fz). The response time course is averaged across all repetitions of the rhythmic pattern making up the stimulus sequences. Note that the auditory response time course shows transient responses to single sensory events composing the pattern (in grey), embedded into slow fluctuations of the signal. In contrast, somatosensory responses do not exhibit such slow fluctuations. On the right, the grand-average magnitude spectra show magnitude of the auditory response located within 0-15 Hz and mainly below 5 Hz, and magnitude of the somatosensory response spread over 0-25 Hz (beat-related and - unrelated frequencies in red and blue respectively), in line with the estimated modality-specific EEG response frequency bandwidths displayed in Fig. 2. Inserts show scalp topographies of the averaged magnitude at beat-related and -unrelated frequencies within each modality-specific frequency bandwidth.

**Figure 6.**
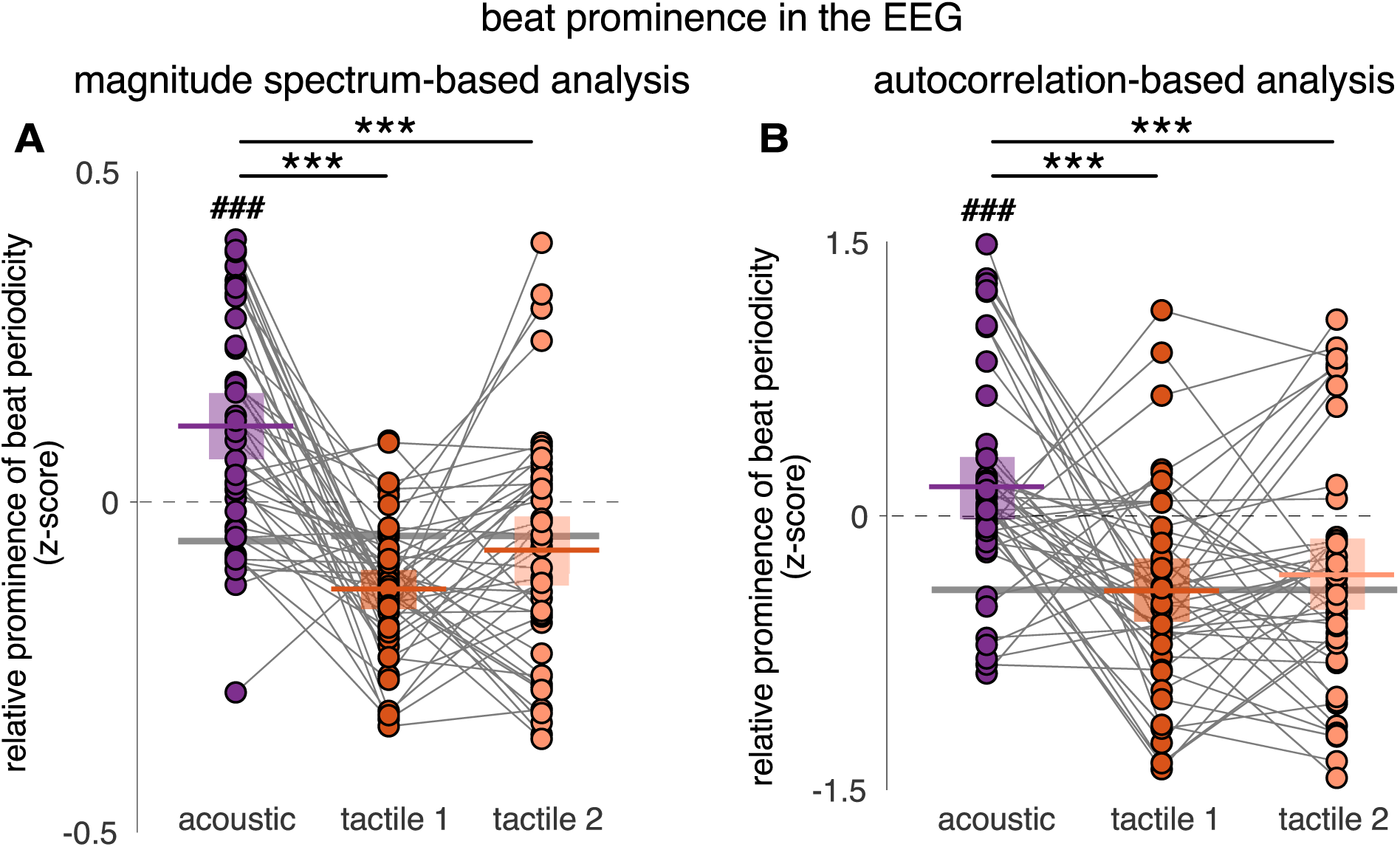
EEG responses show greater prominence of the beat periodicity for acoustic vs. tactile rhythm, in line with tapping responses. **A.** Magnitude spectrum-based analysis. **B.** Autocorrelation-based analysis. Note the significantly reduced periodization for tactile vs. acoustic inputs (cross-block and against-stimulus comparison). Horizontal bars represent means and boxes indicate 95% confidence intervals. Each dot represent a participant. The horizontal grey bars correspond to the stimuli z-scored magnitude at beat-related frequencies (**A**) or lags (**B**). Asterisks indicate significant differences between blocks obtained from the post-hoc pairwise comparisons (***p < 0.001). The octothorpe indicate significant differences obtained from the one-sided one-sample t-tests of the EEG z-scores against the stimulus z-scores (###p < 0.001).

In contrast to the acoustic rhythm, the tactile rhythm produces responses over a wider frequency range, with responses concentrated at 5 Hz (1/0.2 s, i.e., the shortest inter-onset interval) and harmonics up to 25 Hz. In the time domain, this higher-frequency activity takes the form of short transient responses faithfully tracking each event onset and returning to baseline before the next onset occurs. Such a faster time-scale is in line with the preferential response bandwidth reported for somatosensory evoked responses of 20-30 Hz (Tobimatsu et al. 1999; Vlaar et al. 2015; Ahn et al. 2016). More generally, such a faster response could reflect a more discrete processing of incoming tactile inputs (de Haan and Dijkman 2020), to the detriment of temporal integration over longer timescales.

### Capturing neural activity compatible with primary auditory and somatosensory cortices

The current study captured neural responses to rhythm with topographical distributions compatible with activity originating from the primary auditory (A1) and primary somatosensory (S1) cortices, respectively. More specifically, the scalp topography of the somatosensory response is highly indicative of cortical generators with a dipole axis tangential to the scalp and perpendicular to the central sulcus in the hemisphere contralateral to the stimulated hand, i.e., S1 generators (Allison et al. 1991; Moungou et al. 2016).

The fronto-central topographical distribution of the auditory response is known to mainly reflect activity originating from bilateral Heschl’s gyri, i.e. A1 generators (Picton 2011; Pantev et al. 1988; Tan et al. 2016). However, such a topography does not itself rule out substantial contributions of other median brain regions (Mouraux and Iannetti 2009; Somervail et al. 2020). Nevertheless, based on recent evidence for significant neural emphasis of the beat observed in the human Heschl’s gyrus (Nozaradan et al., 2016; Lenc et al., 2024b), it can be reasonably assumed that A1 is embedded into a brain network enabling this higher-level temporal integration and ultimately yielding beat-related periodization of rhythmic input.

Similarly, S1 is also embedded in a functional network which comprises higher-order associative and motor areas such as the cingulate cortex and supplementary motor area (SMA), as evidenced by studies assessing temporal integration of tactile input in the context of working memory tasks (Harris et al. 2002; Numminen et al. 2004). Importantly, these studies revealed a gradient of temporal integration from S1 to these higher-order brain regions, with the latter specifically engaged into slower, supra-second scales while the former would be crucial for retaining information in the sub-second range. In the present study, the somatosensory response to the tactile rhythm is compatible with activity predominantly originating from S1, which might thus explain the relative lack of temporal integration at longer timescales and associated reduced ability to move to the beat compared to the auditory modality.

### Multisensorial redundancy in rhythm perception: somatosensation as a special case?

When parameters of rhythmic stimuli are tuned to match the sensitivity of the sensory modality, acoustic or visual rhythms have reported to elicit equally good synchronization performances. As shown here, this is not the case for tactile rhythmic stimuli. In other words, there seems to be a redundancy for rhythm between the auditory and visual modalities but not with the somatosensory modality. This difference might be partially explained by the fact that both audition and vision share the property of being possibly stimulated by external inputs from a large distance from the body and from a remote region of the peri-personal space (Macaluso and Maravita 2010; Canzoneri et al., 2012). The auditory and visual modalities are thus more likely stimulated concomitantly by a single external source of rhythmic inputs, particularly when positioned at distance from the body, leading to higher probability for redundancy between the auditory and visual modalities in processing rhythmic inputs.

In contrast, the somatosensory modality requires closer proximity or even direct skin contact with the stimuli for humans to perceive them. In addition, and in contrast to audition and vision, somatosensation and associated functions (such as body perception and ownership) induced and fed by a number of afferent subsystems (tactile, proprioceptive, interoceptive, vestibular; see De Haan and Dijkerman 2020). The integration between these various subsystems, with their distinct receptors and neuroanatomical pathways (Hollins, 2010; Proske and Gandevia, 2012; Saal and Bensmaia 2014), might thus be key to elicit higher-level perceptual experience such as the beat.

Along this line, it could be speculated that loud low-frequency sounds and concomitantly produced vibrations could constitute a functionally relevant input to this multi-entry system (Hove et al. 2020; Cameron et al., 2022). Future research should also try to clarify the extent to which individual experience either due to long-term music practice or sensory loss (i.e., deafness) might shape the ability of the somatosensory system to take over multiscale temporal integration, thus addressing unresolved questions relative to multisensory interaction, learning and brain plasticity in humans.

## Authors contributions

C.L. and S.N. designed the study, C.L. performed the study, C.L. and S.N. analyzed data, created the figures and wrote the paper. All authors contributed to writing and editing the paper.

